# The structure and diversity of strain level variation in vaginal bacteria

**DOI:** 10.1101/2020.06.26.173922

**Authors:** Brett A. Tortelli, Amanda L. Lewis, Justin C. Fay

## Abstract

The vaginal microbiome plays an important role in human health and species of vaginal bacteria have been associated with reproductive disease. Strain level variation is also thought to be important, but the diversity, structure and evolutionary history of vaginal strains is not as well characterized. We developed and validated an approach to measure strain variation from metagenomic data based on SNPs within the core-genomes for six species of vaginal bacteria: *G. vaginalis*, *L. crispatus*, *L. iners*, *L. jensenii*, *L. gasseri*, and *A. vaginae*. Despite inhabiting the same environment, strain diversity and structure varies across species. All species except *L. iners* are characterized by multiple distinct groups of strains. Even so, strain diversity is lower in the *Lactobacillus* species, consistent with a more recent colonization of the human vaginal microbiome. Both strain diversity and the frequency of multi-strain samples is related to species-level diversity of the microbiome in which they occur, suggesting similar ecological factors influencing diversity within the vaginal niche. We conclude that the structure of strain level variation provides both the motivation and means of testing whether strain level differences contribute to the function and health consequences of the vaginal microbiome.

**Data Summary:** All vaginal metagenomic sequence data generated for this project can be found on the Sequence Read Archive under BioProject PRJNA639592.

## Introduction

A diverse range of microbial communities have been found to be associated with human anatomical sites, including the skin, gastrointestinal tract, oral cavity and vagina (1). Surveys of these microbial communities have demonstrated significant differences between anatomical sites but also variation among individuals (1, 2). Inter-individual variation in microbial communities has in many instances been associated with a variety of host factors including human health and disease, e.g. obesity and inflammatory bowel disease (3), leading to continued investigation of the implications of microbial variation.

Inter-individual variation has largely been explored by means of characterizing differences in species presence or relative abundance. However, prevalent species not only show differences in relative abundance but also exhibit appreciable strain level variation (1, 2). Individual strains may be unique to a person’s microbiome and bacterial strains of a species isolated from the same individual have been noted to be more similar to each other than strains isolated from different individuals (2, 4). When examined, strain level variation is characterized by functional differences, prominent among these are differences in metabolic potential and antibiotic resistance (2, 5, 6). This suggests that strain level variation may contribute to phenotypic differences in personal microbiomes observed between individuals. However, knowing the extent to which strain-level differences translate to functionally distinct strains remains an open and important question. Currently, most comparisons of microbial communities utilize operational taxonomic units as means of grouping similar strains together and differentiating them from other groups.

Strain differences and their relationships define the population structure of a species. Population structure is relevant for both grouping strains but also making inferences about their history. In the absence of recombination, strains continually diversify, but those lineages that are most successful will expand and others will be lost. Eventually such lineages can diverge in function and even establish new species. Under the ecological species concept, two species can’t stably coexist unless they differ in their niche (7). However, population structure can also be established by limited migration, in which case subpopulations may have the same functions in the community but diverge (neutrally) in their genome (8). Although distinguishing functional populations from neutral populations is difficult, population structure remains an important component of describing groups of strains with shared functional differences or shared population history. As each human may carry or enable the formation of unique microbial strains, the characterization of population structures and their determinants is important to addressing the role of strain level variation in the human microbiome (9).

Among human microbial communities, the vaginal microbiome differs in its community composition. Both 16S ribosomal profiling and metagenomic community profiling have indicated that the vaginal microbiome often exhibits lower community diversity when compared to other anatomical sites, frequently being dominated by a single species (1, 2). The composition of the bacterial community is often described in terms of five common community types (10). Four of the five community types are dominated by a single *Lactobacillus* species (*L. crispatus*, *L. iners*, *L. jensenii* or *L. gasseri*). The fifth community type is characterized by a lack of *Lactobacillus* dominance and often exhibits higher community diversity. This diverse community has been correlated with a high vaginal pH (> 4.5) and bacterial vaginosis (BV) (10), a dysbiosis associated with the overgrowth of anaerobic bacteria including *G. vaginalis* and *A. vaginae*. The prevalence of these vaginal communities vary by self-reported race/ethnicity (10, 11) and have been associated with reproductive health (12). While community type classification offers a convenient method for categorizing the overall composition of the vaginal microbiome, the significance of strain level variation is of increasing interest.

For certain vaginal bacterial species, functional, phenotypic and genomic differences have been described among isolated strains. An example of this is the classification of *G. vaginalis* into distinct phylogenetic clades (groups) through genomic approaches such as gene ontology and genome-wide single-nucleotide polymorphism (SNP) analysis (13–15). Characterization of individual strains have shown functional differences (including sialidase activity and vaginolysin production) between groups with phenotypic consequences (16–19). Such functional differences may explain why some groups but not others have been associated with BV (20–22). While less is known about other vaginal bacterial species, genomic analysis of *L. crispatus* and *L. iners* strains has provided some insight into the population structure of these species (4, 23, 24). An examination of 41 strains found that *L. crispatus* may be comprised of two closely related groups (4), but identification of phenotypic differences between these groups is lacking. Additionally, the population structure of *L. iners* appears to lack strain groupings, but rather each strain appears to be distinct (4, 24). These assessments of strain level variation have focused on isolated strains, and assessments of strain level variation within the vaginal microbiome have been limited (4). The use of variable regions of the 16S gene to define genovariants has been used by some as a proxy for strain diversity (4, 25, 26). However, the use of 16S genovariants to explore strain level variation and associations with health is limited by the resolution of genovariants and their correspondence to phylogeny.

A critical factor in evaluating strain level variation is how it is measured. Early studies employed multilocus sequence typing (MLST) (27), but recombination and horizontal gene transfer (HGT) can cause results to differ depending on the loci employed. Strain level variation has also been examined using gene ontology or copy number variation (CNV) analysis (4, 6), which has the advantage that many CNVs are functionally important. However, CNV can be hard to detect in low coverage samples and HGT can stimulate CNV (28). Genome-wide SNP analysis has also been used (2, 4, 5), but limited reference genomes for some species and variation in genome content present challenges (29). Furthermore, widely divergent species may have limited core genomes and alignment methods for such divergent species present significant difficulties (30). Additionally, it can be challenging to distinguish between strains with mixed ancestry from multi-strain samples.

The goal of the present study was to define and compare the population structure of common vaginal bacteria and identify patterns of strain level variation among vaginal microbiomes. We developed and validated a genome-wide SNP analysis based on available reference genomes. We applied this approach to metagenomic data from vaginal samples and found that diversity present among the vaginal samples was well represented by the available reference strains. We found species-specific differences in strain variation and structure, identifying clear groupings within most of the species. Although our power was limited, no strong associations between strain and host factors were identified. Together, our results provide insight into how vaginal microbiome community types developed over the course of human history and lay the groundwork for assessing the importance of strain level variation in the vaginal microbiome and human health.

## Methods

### Metagenomic sequencing of vaginal samples

We obtained 197 cervicovaginal swabs from 195 pregnant women: 25 cervical swabs and 142 vaginal swabs (collectively referred to as vaginal samples) through the Global Alliance to Prevent Prematurity and Stillbirth (GAPPS) biobank and 30 vaginal swabs from the Women and Infants Health Consortium (WIHSC) at Washington University in St. Louis (IRB #201610121). When selecting samples from the GAPPS biobank, efforts were made to: 1) select all available specimens from women who delivered preterm (< 37 weeks of gestational age), 2) increase the representation of specimens from women of non-White race/ethnicity among the cohort, and 3) balance samples across all 3 trimesters. We augmented the samples selected from GAPPS with samples obtained from women currently enrolled in other studies with WIHSC. Patient data including gestational age at time of swab collection, gestational age at the time of birth, birthweight, maternal age and race/ethnicity were obtained from GAPPS and WIHSC. Women who delivered prior to 37 weeks of gestational age were considered preterm and represented both spontaneous and indicated preterm delivery. To extract genomic DNA, frozen vaginal swabs were eluted in 250 μL of an enzyme solution containing 0.5 mg ml^−1^ lysosozyme, 150 units ml^−1^ mutanolysin, 12 units ml^−1^ lysostaphin, 0.025 units ml^−1^ zymolase in 0.05 M potassium phosphate buffer (pH 7.5) and incubated for 1 hour at 37°C. A ZR Fungal/Bacterial DNA MicroPrep Kit (Zymo Research) was used to extract and purify genomic DNA from swab elutions. Metagenomic sequencing libraries were prepared with a Nextera DNA Sample Prep Kit (Illumina) using 5 ng of genomic DNA and a small volume protocol (31). PCR was performed using KAPA Hi-Fi HotStart ReadyMix (KAPA Biosystems) and libraries were purified with AMPure XP magnetic beads (Beckman Coulter). Libraries were pooled and sequenced on an Illumina NextSeq platform (75 cycles).

### Sequence processing and classification of the vaginal microbiome

Sequence reads from our vaginal samples were trimmed and quality filtered using fqtrim (version 0.9.7) to remove reads less than 50 basepairs in length and trim read ends where quality scores drop below 10. Reads were then aligned to the human genome with Bowtie2 (32) (version 2.3.4) and human reads were discarded. Metagenomic data from 128 vaginal specimens collected as part of the Human Microbiome Project (1) were obtained from NCBI’s Sequence Read Archive (SRP002163). Data were filtered to remove reads less than 50 basepairs in length and remove human reads comprised of Ns using fastq-mcf (version 1.04.803). Taxonomic profiling was performed on non-human reads using MetaPhlAn2 (33) (version 2.6.0). Each microbiome was classified into community types based on the dominant *Lactobacillus* species present, defined as 50% relative abundance or greater and referred to as, “*L. crispatus-*dominant”, “*L. iners-*dominant”, “*L. gasseri-*dominant”, or “*L. jensenii*-dominant”. Communities without a single *Lactobacillus* species reaching 50% were referred to as “diverse” communities. Read data for all vaginal samples were aligned with BWA and Stampy as described below.

### Description of our clinical cohort

For the 195 women in our study, we obtain clinical and demographic data. Data on self-reported race/ethnicity showed most (96%) reported White, Black, Hispanic or Asian. The remaining women (4%) reported either American Indian/Alaskan Native, multiple races or their race/ethnicity was unknown. Maternal age (years), gestational age at sample collection (days), birthweight (grams) and gestational age at delivery (days) was also collected. Preterm delivery was defined as delivery prior to 37 weeks. Sixty-nine (35%) women had *L. crispatus*-dominant microbiomes, 53 (27%) had *L. iners*-dominant microbiomes, 9 (5%) had *L. jensenii*-dominant microbiomes, 9 (5%) had *L. gasseri*-dominant microbiomes and 55 (28%) had diverse microbiomes. A summary of clinical and demographic data can be found in Table S1.We noted a higher prevalence of *L. crispatus*-dominant microbiomes among White (42%) than Hispanic (32%) or Black (20%) women; a higher prevalence of *L. iners*-dominant microbiomes among Hispanic (39%) and Black (35%) than White women (19%); and a higher prevalence of diverse microbiomes among Black (45%) than White (25%) or Hispanic (25%) women (Table S2).

### Reference strain analysis and validation

As reference for the vaginal samples and to identify the core genome we obtained genome assemblies for reference strains from NCBI for the six bacterial species of interest. A total of 101 *G. vaginalis*, 60 *L. crispatus*, 21 *L. iners*, 18 *L. jensenii*, 31 *L. gasseri*, and 5 *A. vaginae* assemblies were obtained (Table S3). Two *G. vaginalis* strains were not included in our analysis: UMB0388 which mapped extensively to other genomes, suggesting the assembly was not a pure isolate; and 6420LIT which had a particularly small genome when compared to all other *G. vaginalis* genomes. ART-MountRainer (34) (version 2.5.8) was used to generate simulated Illumina data (75 basepair reads, NextSeq500 platform v2) at 20x coverage for each assembly.

Simulated read data were aligned to a concatenated reference database containing a representative assembly for each species (Table S4). A concatenated database was used in order to eliminate reads with low mapping quality due to equivalent mappings to multiple species. Alignments were performed with Stampy (30) (version 1.0.32) using the BWA-facilitated option and with an expected divergence of 0.05. Alignment data was filtered to remove reads with a mapping quality score of less than 10 using Samtools (35) (version 1.9).

We identified single nucleotide polymorphisms (SNPs) among all reference strains for a species and removed all variants not present within the core genome. The core genome was defined as all sites that had coverage for all reference strains in each species, and represented 12% (*G. vaginalis*), 47% (*L. crispatus*), 82% (*L. iners*), 72% (*L. jensenii*), 72% (*L. gasseri*) and 24% (*A. vaginae*) of the genome. Single nucleotide polymorphisms (SNPs) were called using GATK UnifiedGenotyper (36, 37) (version 3.6.0). Variant calls were filtered using GATK to remove variants that met the following criteria: QD < 50, FS > 60.0, MQ < 40.0, MQRankSum < −12.5, and ReadPosRankSum < −8.0. A small number of variants were removed due to cross-species read mapping. These sites were called based on reads from one species mapping to an incorrect reference genome, e.g. variant calls in the *L. crispatus* genome based on simulated reads from an *L. iners* reference assembly. Variant selection and removal was completed using VCFtools (38) (version 0.1.14). Among the reference strains, variant sites represented 15% (*G. vaginalis*), 3% (*L. crispatus*), 3% (*L. iners*), 2% (*L. jensenii*), 4% (*L. gasseri*) and 17% (*A. vaginae*) of the core genome.

To determine whether the choice of reference genome for read mapping affected the relationship between reference strains we aligned all simulated reference strain data to a second set of alternative reference genomes (Table S4) and genotyped variants as described above. Variant sites with more than 2 alleles were then removed from both the original and alternate call set using VCFtools (38) (version 0.1.14) and genotypes were extracted using a custom script. A Euclidean distance matrix of strains for each species and each mapping reference was generated and compared by a Mantel test in R (version 3.5.1). The distance matrices for each mapping reference were found to be significantly correlated (Pearson’s correlation coefficient > 0.9, p < 0.05) for each species, indicating that the choice of mapping reference genome did not alter the relationships among strains. Based on this finding we utilized a single mapping reference set (Primary Reference Set) for all analyses.

### Alignment and SNP calling for vaginal samples

Sequence reads from vaginal samples obtained for this study (N = 197) and HMP samples (N = 128) were aligned to the reference genomes as described above. SNPs were independently called from alignment files for all 325 vaginal samples and 233 reference strains and filtered using GATK as described above. SNPs outside of the core genome were removed. Genotype calls with less than 4x coverage were removed with VCFtools.

### Population structure

The core genome variants were filtered to select for biallelic sites with VCFtools. We then removed vaginal samples with > 50% missing sites. The set of samples that passed this filter were considered to have adequate coverage for variant analysis of that species. Variant sites with > 10% missing genotypes across all samples were removed. Heatmaps of the hierarchically clustered variants and samples were generated in R using the heatmap function. Principal component analysis (PCA) was performed on the variant data in R with the package ‘FactoMineR’. All principal components (PCs) explaining > 5% variance were assessed for associations with host factors (see Statistical Methods). For *L. iners* where no PC explained > 5% of the variance, we assessed the PC that explained the most variance.

VCF files containing core genome SNPs for reference strains and vaginal samples were converted to PLINK format with PLINK (39) (version 1.9). ADMIXTURE (40, 41) (version 1.3.0) was then used to identify populations and infer ancestry. The number of populations (groups) was estimated based on the cross validation (CV) error for the number of groups {K1…10}. The estimated number of groups (K) was identified as the point at which the CV error plateaued to a minimum. Reference strains and samples with < 90% of ancestry estimated to be derived from a single group were classified as mixed ancestry (Table S5).

For *G. vaginalis* and *L. crispatus* we estimated group abundance within vaginal samples using the relative allele depth at group-specific SNPs. Group-specific SNPs were defined as those with an allele frequency of 80% or more in one group but 20% or less in all other groups. Based on this designation we identified 22 (Group 1), 210 (Group 2), 19 (Group 3) and 178 (Group 4) SNPs out of 4,884 SNPs in the 88 G. vaginalis reference strains with less than 10% mixed ancestry. For *L. crispatus* we identified 604 (Group 1), 55 (Group 2) and 75 (Group 3) SNPs out of 4,469 SNPs. Using these SNPs we extracted the allele depth supporting each nominally heterozygous genotype. The abundance of each group in each mixed sample was estimated by the average proportion of allele depth for each group-specific SNP.

### Phylogenetic Analysis

Reference strains without mixed ancestry were used for phylogenetic analysis. Four-fold degenerate synonymous SNPs were selected using SNPeff (42) (version 4.3T) and SNPsift (43) (version 4.0). The SNPs for each sample were concatenated into FASTA format with VCF-kit (44). The number of 4-fold synonymous sites surveyed was determined by identifying all 4-fold synonymous sites within the core genome using the same filters as described above except both variant and non-variant sites were retained. A distance matrix of pairwise differences in 4-fold synonymous SNPs/4-fold synonymous sites surveyed (4π) was used to generate a neighbor joining (NJ) tree in R with the packages ‘ape’ and ‘phangorn’. We calculated Watterson’s estimator of diversity (45) with the formula: 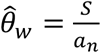, where S= the number of SNPs, 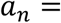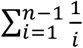 and n is the sample size.

In parallel we identified 0-fold degenerate non-synonymous SNPs and 0-fold non-synonymous sites surveyed. The average pairwise difference in 0-fold non-synonymous variant sites/0-fold non-synonymous sites surveyed (0π) were determined. Tajima’s D was calculated as previously described (46).

For each species we estimated the average time to the most recent common ancestor (TMRCA) in generations as t=d/(2μ), where μ is the mutation rate and d is the average or maximum pairwise distance between strains at synonymous sites. We used a bacterial mutation rate of 2e^−10^ mutations per base pair per replication from *E. coli* (47). We used an *in vitro* doubling rate of *G. vaginalis* (7.1 hours) (48) to estimate a replication rate of approximately 3.38 generations per day for all of the species and convert time in generations to time in years.

### Statistical analysis

All principal components (PCs) explaining > 5% variance were assessed for associations with host factors. For *L. iners* where no PC explained > 5% of the variance, we assessed the PC that explained the most variance. Kruskal-Wallis and Spearman rank correlation tests were used to test for associations between PCs and host factors as appropriate. Due to multiple comparisons, a p-value below 0.001 was considered significant for associations with principal components and a p-value below 0.05 was considered significant for associations with microbiome community type. Statistical analysis was conducted in R.

## Results

To characterize the diversity and structure of variation within common vaginal bacterial species, we generated metagenomic data from 197 vaginal swabs. A median of 3.37e^6^ reads per sample remained after removing human reads, providing adequate coverage of the microbial genome. An analysis of the composition of the microbiome using MetaPhlAn2 (33) indicated compositions similar to those described in prior studies with *Lactobacillus*-dominant and diverse community types: 71 (36.0%) were *L. crispatus*-dominant, 9 (4.6%) were *L. jensenii*-dominant, 53 (26.9%) were *L. iners*-dominant, 55 (27.9%) were diverse and 9 (4.6%) were *L. gasseri*-dominant (Table S1). The prevalence of these community types differed by self-reported race/ethnicity (Table S2).

Using MetaPhlAn2 community composition data, we identified species for strain analysis that were well represented among our vaginal samples. Six bacterial species (*L. crispatus*, *L. iners*, *L. jensenii*, *L. gasseri*, *G. vaginalis* and *A. vaginae*) showed a minimum relative abundance of 10% in at least 10 samples. When metagenomic data was mapped to a set of reference genomes (Table S4), most of the samples (60%) showed a minimum of 4x coverage to one or more of the bacterial species. This indicated that most of the samples had sufficient shotgun metagenomic data to identify single nucleotide polymorphisms (SNPs) and examine strain level variation. To increase the number of metagenomic samples for strain analysis, we included data from an additional 128 vaginal samples collected as part of the Human Microbiome Project (HMP) (1). A summary of the number of metagenomic samples included in the strain analysis is presented in Table 1.

**Table 1.** Metagenomic vaginal samples included in strain analysis for each species.

|  | *G. vaginalis* | *L. crispatus* | *L. iners* | *L. jensenii* | *L. gasseri* | *A. vaginae* |
| --- | --- | --- | --- | --- | --- | --- |
| **Total samples** | 67 (100) | 137 (100) | 115 (100) | 53 (100) | 32 (100) | 30 (100) |
| **From this study** | 42 (62.7) | 71 (51.8) | 70 (60.9) | 20 (37.7) | 16 (50.0) | 16 (53.3) |
| **From HMP** | 25 (37.3) | 66 (48.2) | 45 (39.1) | 33 (62.3) | 16 (50.0) | 14 (46.7) |
| **Multi-strain samples** | 42 (62.7) | 29 (21.2) | 22 (19.1) | 1 (1.9) | 3 (9.4) | 23 (76.7) |

Values are n (%)

A significant challenge to SNP identification from mixed metagenomic samples is being able to reliably call variants for the correct species. This is complicated by horizontal gene transfer, close relationships among the *Lactobacillus* species and variation in genome content. To address these issues, we generated simulated metagenomic read data from publicly available reference genomes (Table S3) for each of the bacterial species (99 *G. vaginalis*, 60 *L. crispatus*, 21 *L. iners*, 18 *L. jensenii*, 31 *L. gasseri* and 5 *A. vaginae*). We aligned the simulated metagenomic reads to the reference set and found low mean misalignment rates for each species: 0.011 (*G. vaginalis*), 0.020 (*L. crispatus*), 0.003 (*L. iners*), 0.018 (*L. jensenii*), 0.017 (*L. gasseri*) and 0.092 (*A. vaginae*). While infrequent, misalignment did result in a small number of SNPs being called to the wrong species (e.g. variant calls in the *L. crispatus* genome based on simulated reads from an *L. iners* reference assembly). These invalid SNPs were excluded from our analysis. Next we identified the core genome SNPs for each species, based on sites represented among all reference strains. Within the core genome, we identified thousands of SNPs for each of the species (Table 2).

**Table 2.** SNP counts and nucleotide diversity measures.

|  | *G. vaginalis* | *L. crispatus* | *L. iners* | *L. jensenii* | *L. gasseri* | *A. vaginae* |
| --- | --- | --- | --- | --- | --- | --- |
| **Core genome sites** | 2.01E+05 | 1.00E+06 | 1.06E+06 | 1.19E+06 | 1.41E+06 | 3.52E+05 |
| **Core genome SNPs** | 3.12E+04 | 2.61E+04 | 3.24E+04 | 1.84E+04 | 6.01E+04 | 6.14E+04 |
| **4-fold degenerate SNPs** | 1.04E+04 | 6.60E+03 | 1.10E+04 | 5.49E+03 | 1.85E+04 | 2.33E+04 |
| **0-fold degenerate SNPs** | 7.33E+03 | 8.36E+03 | 7.08E+03 | 3.04E+03 | 1.23E+04 | 1.70E+04 |
| **Average $\pi_4$** | 0.047 | 0.011 | 0.031 | 0.019 | 0.033 | 0.190 |
| **Average $\pi_0$** | 0.003 | 0.002 | 0.003 | 0.001 | 0.003 | 0.032 |
| **Average $\pi_0/\pi_4$** | 0.063 | 0.172 | 0.100 | 0.068 | 0.077 | 0.167 |
| **$\theta_w$** | 0.068 | 0.012 | 0.026 | 0.011 | 0.028 | 0.206 |
| **Tajima's D** | -1.06 | -0.273 | 0.806 | 2.65 | 0.749 | -0.610 |

To evaluate strain variation among vaginal samples, we independently called SNPs among all vaginal samples and reference strains within core genomic regions. When the data were hierarchically clustered based on SNP profiles, clusters of strains were present for all of the species except *L. iners*. Notably, many of the vaginal samples contained numerous genotype calls with both alleles present (nominally heterozygotes) (Table S5), indicating the presence of multiple strains in a single sample (Fig. 1). We conservatively defined samples as having multiple strains present if more than 10% of the SNPs were called heterozygous. Using this definition, we identified a high proportion of multi-strain samples for *G. vaginalis* (63%) and *A. vaginae* (77%) and a lower proportion (<22%) for the Lactobacillus species (Table 1).

**Fig. 1.**
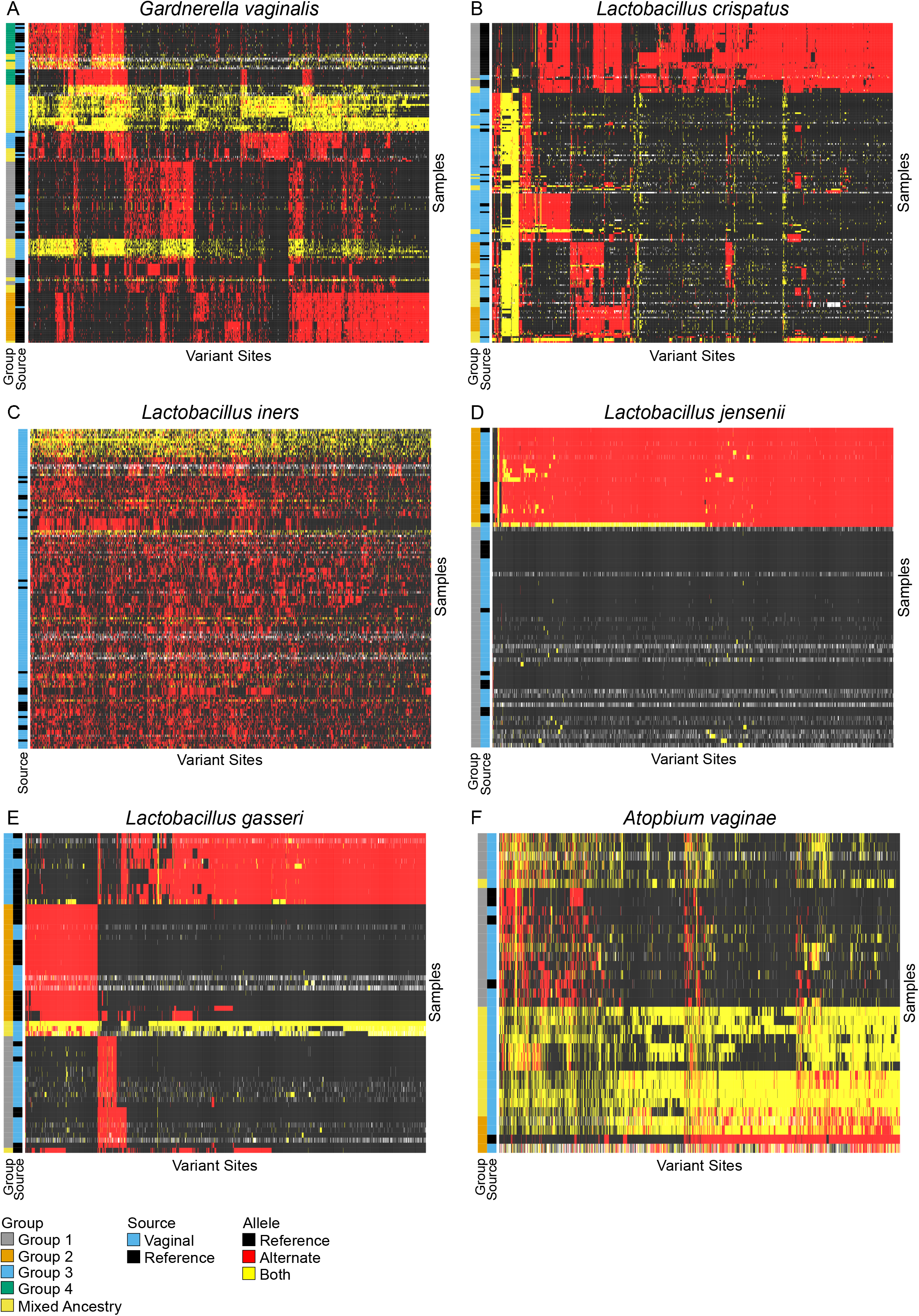
Hierarchically clustered core genome variants for reference and vaginal samples for A) *G. vaginalis*, B) *L. crispatus*, C) *L. iners*, D) *L. jensenii*, E) *L. gasseri* and F) *A. vaginae.* The reference allele is indicated by Black, the alternate allele is indicated by red and the presence of both alleles is indicated by yellow, missing data is indicated by white. To the left of each heatmap is a bar indicating the source of the samples either vaginal or reference strain, and the group to which the strain for each sample was assigned using ADMIXTURE. Samples with mixed ancestry or multi-strain samples from multiple groups are identified as “Mixed Ancestry”.

To compare strain diversity among vaginal samples and isolated reference strains we performed principal component analysis (PCA) of the core genome SNPs. Plotting vaginal samples and reference strains by principal components (PCs) revealed that most reference strains formed clusters and the reference strains represent much of the variation observed among the vaginal samples (Fig. 2).

**Fig. 2.**
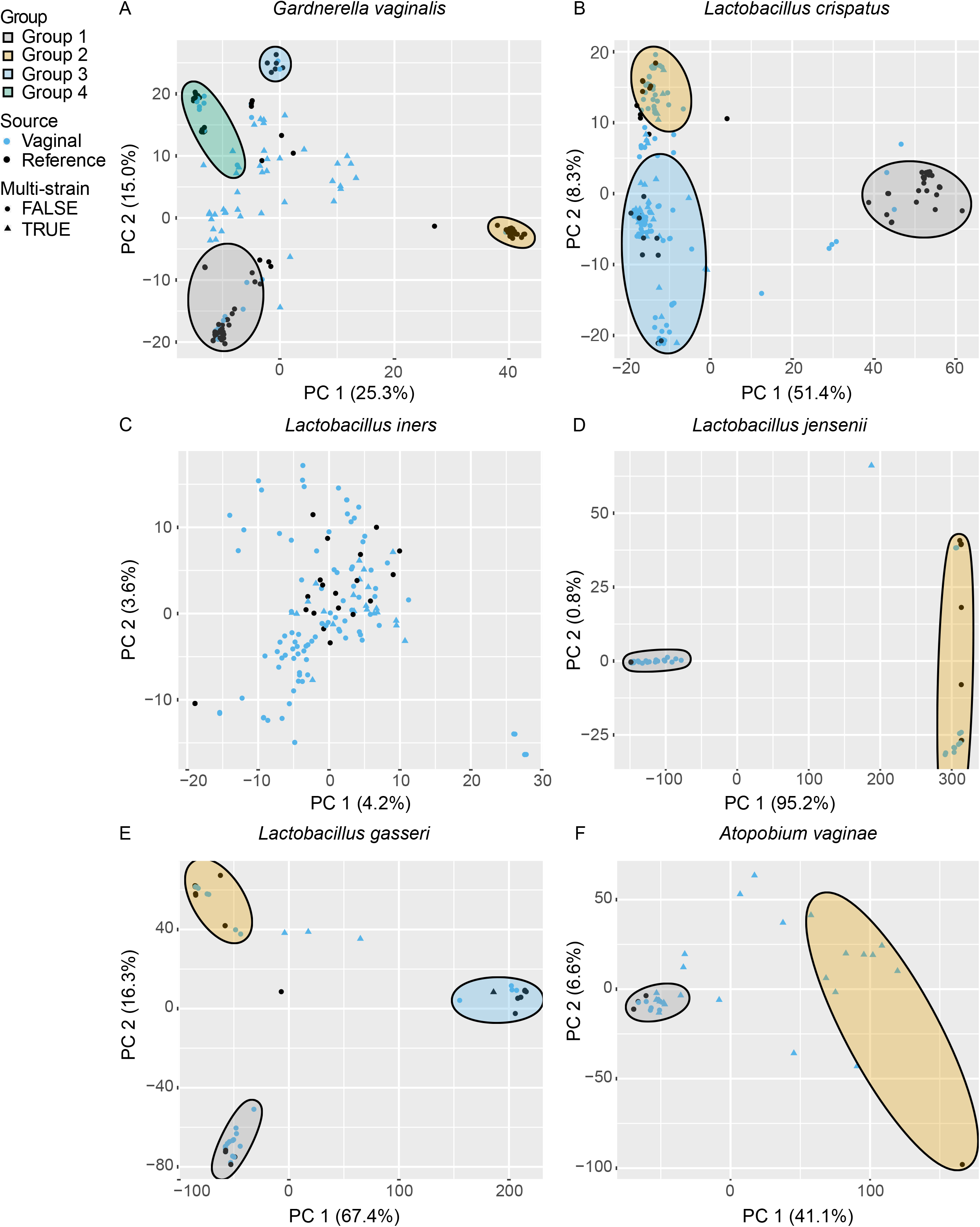
Principal component analysis (PCA) of SNP data for A) *G. vaginalis*, B) *L. crispatus*, C) *L. iners*, D) *L. jensenii*, E) *L. gasseri* and F) *A. vaginae.* Each vaginal sample and reference strain is shown by their PC1 and PC2 coordinates. Vaginal samples are indicated by blue points and reference strains are indicated by black points. Samples that were identified as multi-strain are represented by triangles and single strain samples are represented by circles. Subpopulation groups were determined by ADMIXTURE analysis and ellipses drawn to show vaginal samples and reference strains belonging to a single group.

PCA can distinguish different strain groups but it does not identify strains of mixed ancestry, which can occur through conjugation, transduction and transformation. To examine the structure of strain diversity within each species we used ADMIXTURE (40, 41) to classify samples into groups (subpopulations) and identify samples with mixed ancestry to multiple groups. This analysis identified multiple groups for most of the vaginal species we studied: four *G. vaginalis*, three *L. crispatus*, three *L. gasseri* and two each for *L. jensenii* and *A. vaginae*. We did not identify any population structure for *L. iners*. These groups closely correspond to the clusters observed by PCA (Fig. 2).

Consistent with the high proportion of multi-strain samples, many of the vaginal samples were inferred to be mixtures of groups (Table S5). While ADMIXTURE is unable to distinguish vaginal samples with distinct strains from multiple groups, from samples with a single strain of mixed ancestry, it can identify reference strains with mixed ancestry. Among our reference panel, mixed ancestry was present but uncommon, representing 11 of 99 (11.1%) *G. vaginalis*, 8 of 60 (13.3%) *L. crispatus*, 0 of 18 (0.0%) *L. jensenii*, 1 of 31 (3.2%) *L. gasseri*, and 0 of 5 (0.0%) *A. vaginae* strains (Table S5).

Among the vaginal samples inferred to have mixed ancestry, many were multi-strain samples. In these samples, the relative abundance of reads supporting each allele should be indicative of strain frequency in a sample. We thus used allele-specific read counts of population-specific SNPs to quantify relative abundance of each group in mixed vaginal samples. This was done for *G. vaginalis* and *L. crispatus*, which both have well defined groups based on multiple reference samples and an appreciable number of multi-strain vaginal samples. For each of the multi-strain samples the relative abundance of each group ranged from 10-90% (Table S6), indicating inter-individual variation in strain frequency as well as strain type (Fig. S1). Additionally, the frequencies of different groups are inter-related in *G. vaginalis*. The frequency of Group 4 is negatively correlated with all other groups, whereas among the other groups only Group 1 and Group 3 are negatively correlated with each other (Table S7).

Patterns and levels of strain level variation and population structure are shaped by historical effective population sizes, relationships among groups and strength of selection. To identify the relationships among strains and inferred groups we generated phylogenetic trees for reference strains without mixed ancestry. We constructed phylogenetic trees using synonymous four-fold degenerate sites for 88 *G. vaginalis*, 52 *L. crispatus*, 21 *L. iners*, 18 *L. jensenii*, 30 *L. gasseri* and five *A. vaginae* reference strains (Fig. 3). Groups identified by ADMIXTURE could clearly be identified in the phylogenetic trees (Fig. 3). We then estimated the genetic diversity within each species using the Watterson’s estimator 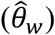 (45). 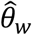 was high for both *G. vaginalis* (0.068) and *A. vaginae* (0.206) reflecting greater genetic diversity when compared to the lower values observed for *L. crispatus* (0.012), *L. iners* (0.026), *L. jensenii* (0.011) and *L. gasseri* (0.028) (Table 2). Tajima’s D measures the relative abundance of common versus rare alleles (46). Population bottlenecks and population structure are expected to generate positive Tajima’s D values and historical expansion of population size is expected to generate negative Tajima’s D values. Tajima’s D was negative for *G. vaginalis* (−1.062), *A. vaginae* (−0.610) and *L. crispatus* (−0.273) and positive for *L. iners* (0.806), *L. jensenii* (2.650) and *L. gasseri* (0.749) (Table 2).

**Fig. 3.**
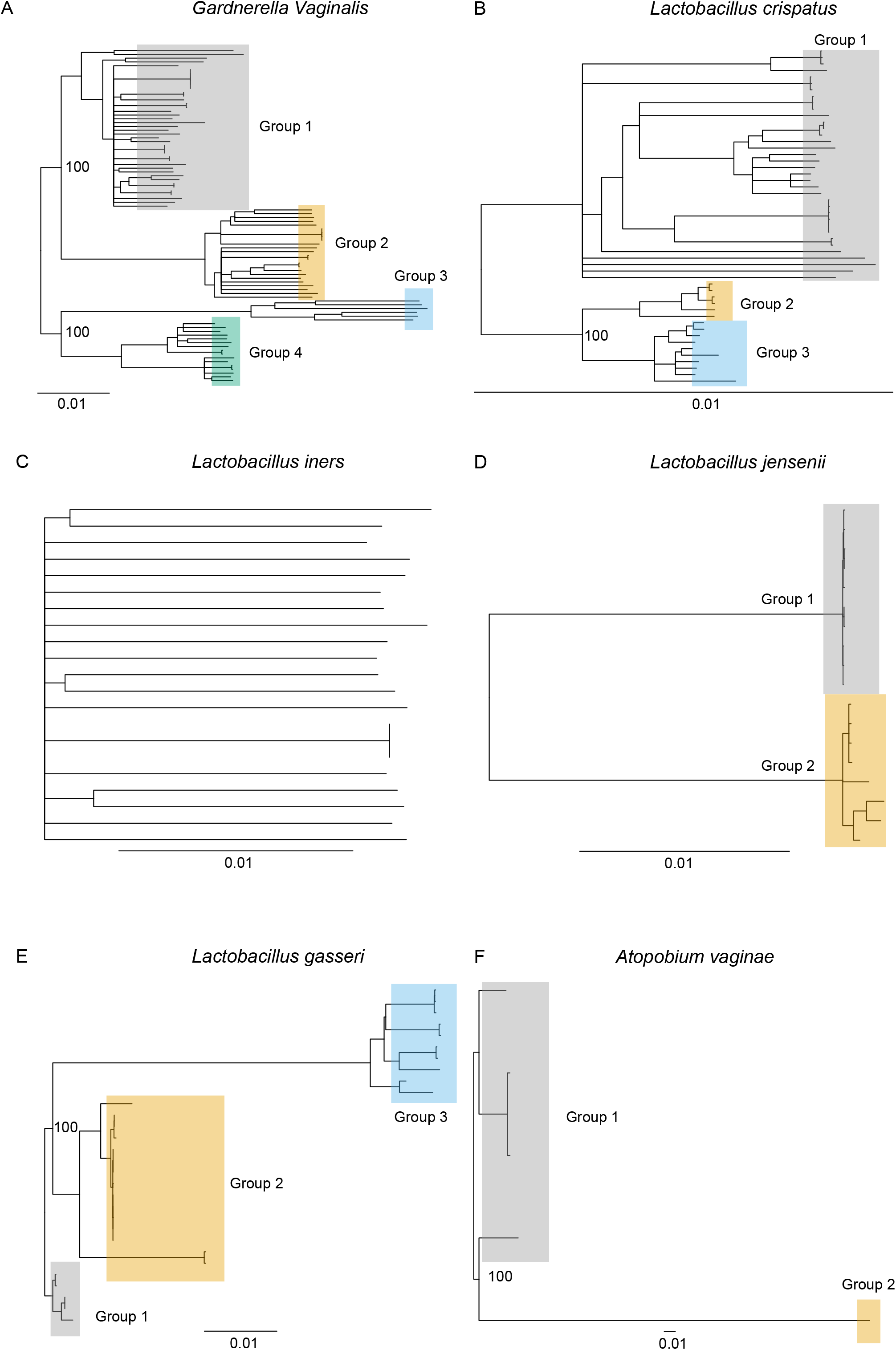
Neighbor joining trees of reference strains created from synonymous sites for A) *G. vaginalis*, B) *L. crispatus*, C) *L. iners*, D) *L. jensenii*, E) *L. gasseri*, F) *A. vaginae*. Branch lengths represent pairwise differences per site surveyed. Groups were determined by ADMIXTURE analysis. Select bootstrap values for nodes separating groups are shown. Bootstrap values indicate the number of supporting iterations out of 100 as calculated by resampling with replacement.

The ratio of nonsynonymous to synonymous diversity is indicative of past selective pressure on a species. Species with higher constraints have lower ratios whereas species with low or altered constraints have higher ratios. We measured diversity at 0-fold degenerate sites (nonsynonymous) and compared it to 4-fold degenerate sites (synonymous). Two of the species, *L. crispatus* and *A. vaginae* had a much higher ratio of 0-fold to 4-fold diversity, 0.172 and 0.167 respectively, compared to the other species (Table 2).

The population structure of vaginal species raises the possibility that subpopulations may exhibit associations with their human host, similar to those associations present at the species level. To test for such associations, principal component values were extracted as a proxy for strain relationships (including groups) for 195 vaginal samples in our study and tested for associations with host factors (race/ethnicity, age and microbiome community type) and birth outcomes (preterm delivery and birth weight) (Table S8). We did not observe any statistically significant correlations (Table S9); however, power analysis suggests that our study was only powered to detect large effect sizes.

## Discussion

Strain level variation is thought to be functional, motivating fine scale measurement of strain variation and testing for its association with reproductive health. In this study we developed and validated a reference genome-based analysis of metagenomic vaginal samples to study the structure of strain level variation within and between individuals. We find reference genomes encompass the majority of strain level vaginal samples, thereby providing a means interpreting strain level variation and structure in vaginal samples. Despite occupying the same environment, we find differences in strain level variation, multi-strain samples, population structure and strength of selection among the vaginal species. Below, we discuss these results in relation to prior studies of strain level variation, and the ecology and evolutionary history of vaginal strains relevant to identifying functional differences among groups and their role in human health.

### Strain level variation

Our analysis of core genomic SNPs provides fine-scale measures of strain level variation and captures known and new aspects of population structure present in vaginal species. Previous genomic studies encompassed only reference genomes, were limited to smaller sample sizes, or did not accommodate multi-strain or admixed samples (Table S10). We made use of combined metagenomic and reference genomes to 1) survey metagenomic SNPs in core genomic regions and establish that most metagenomic variation is captured in reference genomes, and 2) identify groups or subpopulations within each species while accounting for a number of mixed ancestry genomes and numerous multi-strain samples. Mixed ancestry due to recombination or HGT can confound phylogenetic analysis of strain level variation (49), and multi-strain samples are difficult to resolve due to the challenges of accurate assemblies from metagenomic data (50, 51). While our approach does not resolve multi-strain samples into individual lineages, we make use of multiple allele genotype calls to estimate relative abundance of groups within multi-strain samples.

With the exception of strains showing mixed ancestry, the structure of bacterial strain diversity is largely consistent with prior studies (Table S10). We find no population structure of *L. iners*, consistent with previous genomic analyses that showed a highly conserved genome among *L. iners* strains with little difference in gene content (4, 24). A lack of population structure does not convey a lack of strain diversity. Indeed, most *L. iners* strains identified appeared to be unique and average nucleotide diversity among *L. iners* strains was greater than that seen among *L. crispatus* strains (Table 1). The three *L. gasseri* groups we identified correspond to two previously defined groups with distinct gene content (52). Notably, recent studies (53, 54) suggest our *L. gasseri* Group 3 represents the closely related *L. paragasseri*. The three *L. crispatus* groups correlate with two previously reported groups described in a genomic analysis of 41 strains (4). Notably, most vaginal samples harbored Group 2 or Group 3 strains, while reference isolates from avian hosts were common in the more diverse Group 1. This suggests that Group 1 strain colonization of the human vagina may be rare.

The four groups of *G. vaginalis* that we found encompass and are largely consistent with prior groups (Table S11) (4, 13–15). However, a number of these previously defined groups correspond to strains we find to have mixed ancestry. *Ahmed et al.* proposed the division of the species into four groups after a phylogenetic analysis of the core genome of 17 isolated strains (13). Subsequent studies of the *cpn60* gene (17, 22, 55) as well as our strain group assignments are consistent with those described by *Ahmed et al.* (13). One exception is that strains 1400E and 55152, which were assigned to Group 1 but our analysis suggested were of mixed ancestry (mostly Groups 1 and 2). However, assemblies may give the appearance of mixed ancestry if unknowingly generated using a mixture of two or more strains. More recent studies (4, 14, 15) have expanded the number of strains as well as the number of groups (Table S10). However, many of these new groups are comprised of strains our analysis indicates are of mixed ancestry. The placement of strains with mixed ancestry into a separate group is not incorrect; such groups may be functionally distinct. While our approach to inferring population structure does not place mixed ancestry strains into separate groups, our results provide insight into the historical origin of these mixed ancestry groups.

### Ecological diversity of the vaginal microbiome

Vaginal microbial diversity has been correlated with reproductive health and determinants of this diversity are of significant clinical interest. Species diversity within the vaginal niche is determined through ecological interactions within that environment. A key correlate of species diversity is vaginal pH. A vaginal pH less than 4.5 is thought of as healthy and is associated with low diversity, *Lactobacillus*-dominated communities (10, 56). It is believed that through the production of lactic acid, *Lactobacillus* species are able to outcompete other vaginal bacteria and dominate that niche (56). Among *Lactobacillus*-dominated microbiomes, multiple *Lactobacillus* species may be present but a single *Lactobacillus* species usually dominates (10). This suggests that these species may occupy very similar niches within the vagina. According to ecological theory, multiple species cannot occupy the same niche indefinitely and one species will eventually outcompete the others (57). Conversely, a more neutral pH correlates with greater diversity and an abundance of BV-associated anaerobes including *G. vaginalis* and *A. vaginae* (10). These polymicrobial communities support multiple species which may be explained by the theory of resource partitioning in which competing species utilize different subsets of resources to occupy niche divisions within an environment (58).

We find that patterns of strain level variation mimic those of species level diversity. We observed greater strain level diversity within *G. vaginalis* and *A. vaginae*, which are found in more species-diverse communities. Furthermore, most samples with *G. vaginalis* (62.7%) and *A. vaginae* (76.7%) harbor multiple strains from different groups, while multi-strain samples among lactobacilli are much less common (2-21%). The high frequency of multi-strain samples is consistent with prior studies of *G. vaginalis* (4, 20–22) and *A. vaginae* (59). The co-occurrence of different groups can be explained by ecotype theory which suggests that different strains of the same species may occupy the same niche if they function as different ecological species (ecotypes), exploiting different resources (7, 18).

The co-occurrence of differentiated groups within vaginal communities is important for understanding group associations with health. The presence of multiple groups of *G. vaginalis* has been correlated with BV (20, 22). We find that the frequency of Group 4 is negatively correlated with the frequency of all other groups in mixed vaginal samples, potentially indicating that it competitively excludes these groups. In contrast, Group 1, 2 and 3 co-occur but only 1 and 3 are negatively correlated. This is particularly interesting as the co-occurrence of multiple groups of *G. vaginalis* has been correlated with BV (20, 22). Additionally, prior studies have failed to show an associated between Group 4 and BV (20, 22), which may indicate that Group 4 strains are less pathogenic. These findings suggest that mixed group communities may confound *G. vaginalis* group associations with vaginal health and should be accounted for in future models.

While a low pH may enable *Lactobacillus* species to exclude high pH species from the vaginal niche, this does not explain why multiple *Lactobacillus* strains are not observed more frequently in the same sample. If *Lactobacillus* groups represented distinct ecotypes, one would expect groups to co-occur as observed with *G. vaginalis* and *A. vaginae*. One potential explanation is that there has not yet been enough time for *Lactobacillus* strains to diversify and evolve resource partitioning strategies.

### Evolutionary origins of strain level diversity

Strain level diversity is indicative of the species’ demographic history, including past changes in population size, population structure and migration between host microbiomes or other environments. Some insight into the evolutionary origins of the vaginal microbiome may be gleaned by comparing it to microbiome composition of other primates. While vaginal microbial signatures of non-human primates are unique to each species, community compositions more closely resemble the diverse structure associated with *G. vaginalis* and *A. vaginae* (60–62). Only humans are dominated by *Lactobacillus* species (60). While *Lactobacillus* species are closely associated with food and agriculture (63), the species that dominate the human vagina have reduced genome sizes when compared to other *Lactobacillus* species suggesting adaptation to the host environment (64).

Among the species studied here, *A. vaginae* and *G. vaginalis* exhibited the greatest diversity. This diversity may be caused in part by ecotype differentiation. High diversity is also consistent with large, long-term populations of *G. vaginalis* and *A. vaginae* in humans as part of an ancestral state. The *Lactobacillus* species showed much less diversity, which may reflect a smaller historic population size or more recent colonization during human history. Among the *Lactobacillus* species, *L. iners* and *L. gasseri* are the most diverse. The diversity of *L. iners* likely reflects a larger population size, consistent with it being the most prevalent (most frequently detected) of the *Lactobacillus* species (10). *L. gasseri* diversity is also higher than the other *Lactobacillus* species, but this is partly caused by strong divisions between groups, which could predate colonization of the human vaginal microbiome. *L. jensenii* has low diversity and a large positive Tajima’s D, consistent with a recent bottleneck, potentially related to colonization of the vaginal niche.

The origin of these different population structures is unknown but comparing estimates of the time it took for these strains to diverge may provide insight. Using estimates for mutation and replication rates (see Methods) we estimated average TMRCA using average pairwise divergence and the maximum TMRCA using divergence of the two most distantly related strains (Table 3). Both estimates indicated *A. vaginae* and *G. vaginalis* groups diverged much earlier than the *Lactobacillus* species. Of the two most commonly found *Lactobacillus* species, *L. iners* appears to have emerged much earlier than *L. crispatus*. Our rough estimates suggest that *L. crispatus* may have emerged around 20-30 thousand years ago, after the time that it is believed modern *Homo sapiens* settled Europe (65). These observations seem to parallel earlier findings from many research groups showing that vaginal microbiomes with an abundance of *G. vaginalis* and/or *L. iners* are more common in women of African descent, whereas an abundance of *L. crispatus* within the vaginal microbiome is more commonly found in women of European descent (10, 11, 66).

**Table 3.** TMRCA estimates for each species.

|  | *G. vaginalis* | *L. crispatus* | *L. iners* | *L. jensenii* | *L. gasseri* | *A. vaginae* |
| --- | --- | --- | --- | --- | --- | --- |
| **Average pairwise difference at synonymous sites** | 0.047 | 0.011 | 0.031 | 0.019 | 0.033 | 0.190 |
| **Average TMRCA<sup>1</sup>** | 9.50E+04 | 2.23E+04 | 6.24E+04 | 3.77E+04 | 6.78E+04 | 3.84E+05 |
| **Maximum pairwise difference at synonymous sites** | 0.089 | 0.015 | 0.035 | 0.036 | 0.071 | 0.394 |
| **Maximum TMRCA<sup>2</sup>** | 1.80E+05 | 3.13E+04 | 7.11E+04 | 7.24E+04 | 1.44E+05 | 7.98E+05 |

<sup>1</sup>Calculated using the average pairwise difference at synonymous sites<sup>2</sup>Calculated using the maximum pairwise difference at synonymous sites

## Conclusion

Our results show that most species are characterized by multiple distinct groups of strains, and that strain diversity and the frequency of multi-strain samples is related to species-level diversity of the microbiome in which they occur. Future work will need to uncover the ecological variables that impact variation within and between communities at the strain level, and the historical genomic and functional differentiation that lead to extant population structure. Doing so will not only help resolve the role of strain variation in the vaginal microbiome as it relates to reproductive disease, but could also provide insight into the establishment and subsequent changes in community composition as a function of important gynecologic and obstetric events including: sexual development, the menstrual cycle, pregnancy and menopause (67).

---

**Fig. S1.**
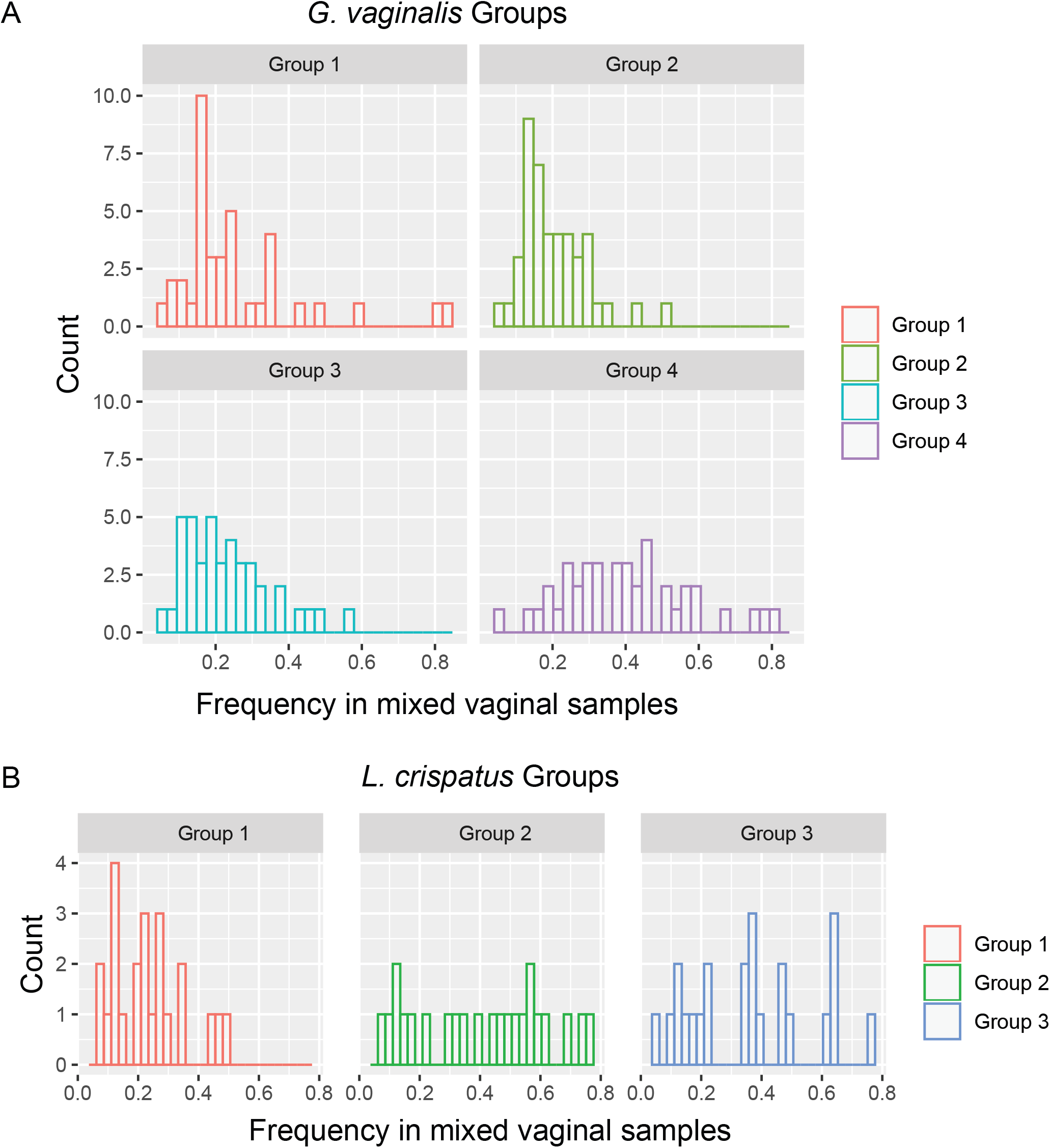
The relative abundance of A) *G. vaginalis* and B) *L. crispatus* groups in mixed samples based on group-specific SNPs. Each panel shows a histogram of the inferred frequency of the four *G. vaginalis* groups and three *L. crispatus* groups in vaginal samples designated as mixed populations by ADMIXTURE.

## References

1. Human Microbiome Project C. Structure, function and diversity of the healthy human microbiome. Nature. 2012;486(7402):207–14.

2. Lloyd-Price J, Mahurkar A, Rahnavard G, Crabtree J, Orvis J, Hall AB, et al. Strains, functions and dynamics in the expanded Human Microbiome Project. Nature. 2017;550(7674):61–6.

3. Gilbert JA, Blaser MJ, Caporaso JG, Jansson JK, Lynch SV, Knight R. Current understanding of the human microbiome. Nat Med. 2018;24(4):392–400.

4. Goltsman DSA, Sun CL, Proctor DM, DiGiulio DB, Robaczewska A, Thomas BC, et al. Metagenomic analysis with strain-level resolution reveals fine-scale variation in the human pregnancy microbiome. Genome Res. 2018;28(10):1467–80.

5. Truong DT, Tett A, Pasolli E, Huttenhower C, Segata N. Microbial strain-level population structure and genetic diversity from metagenomes. Genome Res. 2017;27(4):626–38.

6. Greenblum S, Carr R, Borenstein E. Extensive strain-level copy-number variation across human gut microbiome species. Cell. 2015;160(4):583–94.

7. Cohan FM. Towards a conceptual and operational union of bacterial systematics, ecology, and evolution. Philos Trans R Soc Lond B Biol Sci. 2006;361(1475):1985–96.

8. Sheppard SK, Guttman DS, Fitzgerald JR. Population genomics of bacterial host adaptation. Nat Rev Genet. 2018;19(9):549–65.

9. Schloissnig S, Arumugam M, Sunagawa S, Mitreva M, Tap J, Zhu A, et al. Genomic variation landscape of the human gut microbiome. Nature. 2013;493(7430):45–50.

10. Ravel J, Gajer P, Abdo Z, Schneider GM, Koenig SS, McCulle SL, et al. Vaginal microbiome of reproductive-age women. Proc Natl Acad Sci U S A. 2011;108 Suppl 1:4680–7.

11. Fettweis JM, Brooks JP, Serrano MG, Sheth NU, Girerd PH, Edwards DJ, et al. Differences in vaginal microbiome in African American women versus women of European ancestry. Microbiology. 2014;160(Pt 10):2272–82.

12. Ma B, Forney LJ, Ravel J. Vaginal microbiome: rethinking health and disease. Annu Rev Microbiol. 2012;66:371–371.

13. Ahmed A, Earl J, Retchless A, Hillier SL, Rabe LK, Cherpes TL, et al. Comparative genomic analyses of 17 clinical isolates of Gardnerella vaginalis provide evidence of multiple genetically isolated clades consistent with subspeciation into genovars. J Bacteriol. 2012;194(15):3922–37.

14. Vaneechoutte M, Guschin A, Van Simaey L, Gansemans Y, Van Nieuwerburgh F, Cools P. Emended description of Gardnerella vaginalis and description of Gardnerella leopoldii sp. nov., Gardnerella piotii sp. nov. and Gardnerella swidsinskii sp. nov., with delineation of 13 genomic species within the genus Gardnerella. Int J Syst Evol Microbiol. 2019;69(3):679–87.

15. Potter RF, Burnham CD, Dantas G. In Silico Analysis of Gardnerella Genomospecies Detected in the Setting of Bacterial Vaginosis. Clin Chem. 2019;65(11):1375–87.

16. Li W, Raoult D, Fournier PE. Bacterial strain typing in the genomic era. FEMS Microbiol Rev. 2009;33(5):892–916.

17. Schellenberg JJ, Paramel Jayaprakash T, Withana Gamage N, Patterson MH, Vaneechoutte M, Hill JE. Gardnerella vaginalis Subgroups Defined by cpn60 Sequencing and Sialidase Activity in Isolates from Canada, Belgium and Kenya. PLoS One. 2016;11(1):e0146510.

18. Cornejo OE, Hickey RJ, Suzuki H, Forney LJ. Focusing the diversity of Gardnerella vaginalis through the lens of ecotypes. Evol Appl. 2018;11(3):312–24.

19. Janulaitiene M, Gegzna V, Baranauskiene L, Bulavaite A, Simanavicius M, Pleckaityte M. Phenotypic characterization of Gardnerella vaginalis subgroups suggests differences in their virulence potential. PLoS One. 2018;13(7):e0200625.

20. Janulaitiene M, Paliulyte V, Grinceviciene S, Zakareviciene J, Vladisauskiene A, Marcinkute A, et al. Prevalence and distribution of Gardnerella vaginalis subgroups in women with and without bacterial vaginosis. BMC Infect Dis. 2017;17(1):394.

21. Hill JE, Albert AYK, Group VR. Resolution and Cooccurrence Patterns of Gardnerella leopoldii, G. swidsinskii, G. piotii, and G. vaginalis within the Vaginal Microbiome. Infect Immun. 2019;87(12).

22. Balashov SV, Mordechai E, Adelson ME, Gygax SE. Identification, quantification and subtyping of Gardnerella vaginalis in noncultured clinical vaginal samples by quantitative PCR. J Med Microbiol. 2014;63(Pt 2):162–75.

23. Ojala T, Kankainen M, Castro J, Cerca N, Edelman S, Westerlund-Wikstrom B, et al. Comparative genomics of Lactobacillus crispatus suggests novel mechanisms for the competitive exclusion of Gardnerella vaginalis. BMC Genomics. 2014;15:1070.

24. France MT, Mendes-Soares H, Forney LJ. Genomic Comparisons of Lactobacillus crispatus and Lactobacillus iners Reveal Potential Ecological Drivers of Community Composition in the Vagina. Appl Environ Microbiol. 2016;82(24):7063–73.

25. Callahan BJ, DiGiulio DB, Goltsman DSA, Sun CL, Costello EK, Jeganathan P, et al. Replication and refinement of a vaginal microbial signature of preterm birth in two racially distinct cohorts of US women. Proc Natl Acad Sci U S A. 2017;114(37):9966–71.

26. Callahan BJ, McMurdie PJ, Rosen MJ, Han AW, Johnson AJ, Holmes SP. DADA2: High-resolution sample inference from Illumina amplicon data. Nat Methods. 2016;13(7):581–3.

27. Maiden MC. Multilocus sequence typing of bacteria. Annu Rev Microbiol. 2006;60:561–561.

28. Lind PA, Tobin C, Berg OG, Kurland CG, Andersson DI. Compensatory gene amplification restores fitness after inter-species gene replacements. Mol Microbiol. 2010;75(5):1078–89.

29. Kraal L, Abubucker S, Kota K, Fischbach MA, Mitreva M. The prevalence of species and strains in the human microbiome: a resource for experimental efforts. PLoS One. 2014;9(5):e97279.

30. Lunter G, Goodson M. Stampy: a statistical algorithm for sensitive and fast mapping of Illumina sequence reads. Genome Res. 2011;21(6):936–9.

31. Baym M, Kryazhimskiy S, Lieberman TD, Chung H, Desai MM, Kishony R. Inexpensive multiplexed library preparation for megabase-sized genomes. PLoS One. 2015;10(5):e0128036.

32. Langmead B, Salzberg SL. Fast gapped-read alignment with Bowtie 2. Nat Methods. 2012;9(4):357–9.

33. Truong DT, Franzosa EA, Tickle TL, Scholz M, Weingart G, Pasolli E, et al. MetaPhlAn2 for enhanced metagenomic taxonomic profiling. Nat Methods. 2015;12(10):902–3.

34. Huang W, Li L, Myers JR, Marth GT. ART: a next-generation sequencing read simulator. Bioinformatics. 2012;28(4):593–4.

35. Li H, Handsaker B, Wysoker A, Fennell T, Ruan J, Homer N, et al. The Sequence Alignment/Map format and SAMtools. Bioinformatics. 2009;25(16):2078–9.

36. McKenna A, Hanna M, Banks E, Sivachenko A, Cibulskis K, Kernytsky A, et al. The Genome Analysis Toolkit: a MapReduce framework for analyzing next-generation DNA sequencing data. Genome Res. 2010;20(9):1297–303.

37. DePristo MA, Banks E, Poplin R, Garimella KV, Maguire JR, Hartl C, et al. A framework for variation discovery and genotyping using next-generation DNA sequencing data. Nat Genet. 2011;43(5):491–8.

38. Danecek P, Auton A, Abecasis G, Albers CA, Banks E, DePristo MA, et al. The variant call format and VCFtools. Bioinformatics. 2011;27(15):2156–8.

39. Purcell S, Neale B, Todd-Brown K, Thomas L, Ferreira MA, Bender D, et al. PLINK: a tool set for whole-genome association and population-based linkage analyses. Am J Hum Genet. 2007;81(3):559–75.

40. Palomino HM, Peredo MT, Montenegro MA. [Maternal influence on susceptibility to cleft palate (1)]. Rev Fac Odontol Univ Chile. 1990;8(1):31–6.

41. Alexander DH, Novembre J, Lange K. Fast model-based estimation of ancestry in unrelated individuals. Genome Res. 2009;19(9):1655–64.

42. Cingolani P, Platts A, Wang le L, Coon M, Nguyen T, Wang L, et al. A program for annotating and predicting the effects of single nucleotide polymorphisms, SnpEff: SNPs in the genome of Drosophila melanogaster strain w1118; iso-2; iso-3. Fly (Austin). 2012;6(2):80–92.

43. Cingolani P, Patel VM, Coon M, Nguyen T, Land SJ, Ruden DM, et al. Using Drosophila melanogaster as a Model for Genotoxic Chemical Mutational Studies with a New Program, SnpSift. Front Genet. 2012;3:35.

44. Cook DE, Andersen EC. VCF-kit: assorted utilities for the variant call format. Bioinformatics. 2017;33(10):1581–2.

45. Watterson GA. On the number of segregating sites in genetical models without recombination. Theor Popul Biol. 1975;7(2):256–76.

46. Tajima F. Statistical method for testing the neutral mutation hypothesis by DNA polymorphism. Genetics. 1989;123(3):585–95.

47. Sung W, Ackerman MS, Dillon MM, Platt TG, Fuqua C, Cooper VS, et al. Evolution of the Insertion-Deletion Mutation Rate Across the Tree of Life. G3 (Bethesda). 2016;6(8):2583–91.

48. Khan S, Voordouw MJ, Hill JE. Competition Among Gardnerella Subgroups From the Human Vaginal Microbiome. Front Cell Infect Microbiol. 2019;9:374.

49. Didelot X, Maiden MC. Impact of recombination on bacterial evolution. Trends Microbiol. 2010;18(7):315–22.

50. Nurk S, Meleshko D, Korobeynikov A, Pevzner PA. metaSPAdes: a new versatile metagenomic assembler. Genome Res. 2017;27(5):824–34.

51. Quince C, Delmont TO, Raguideau S, Alneberg J, Darling AE, Collins G, et al. DESMAN: a new tool for de novo extraction of strains from metagenomes. Genome Biol. 2017;18(1):181.

52. Tada I, Tanizawa Y, Endo A, Tohno M, Arita M. Revealing the genomic differences between two subgroups in Lactobacillus gasseri. Biosci Microbiota Food Health. 2017;36(4):155–9.

53. Putonti C, Akhnoukh V, Anagnostopoulos Z, Bilek M, Colgan J, El Idrissi S, et al. Draft Genome Sequences of Six Lactobacillus gasseri and Three Lactobacillus paragasseri Strains Isolated from the Female Bladder. Microbiol Resour Announc. 2019;8(37).

54. Tanizawa Y, Tada I, Kobayashi H, Endo A, Maeno S, Toyoda A, et al. Lactobacillus paragasseri sp. nov., a sister taxon of Lactobacillus gasseri, based on whole-genome sequence analyses. Int J Syst Evol Microbiol. 2018;68(11):3512–7.

55. Paramel Jayaprakash T, Schellenberg JJ, Hill JE. Resolution and characterization of distinct cpn60-based subgroups of Gardnerella vaginalis in the vaginal microbiota. PLoS One. 2012;7(8):e43009.

56. Tachedjian G, Aldunate M, Bradshaw CS, Cone RA. The role of lactic acid production by probiotic Lactobacillus species in vaginal health. Res Microbiol. 2017;168(9-10):782–92.

57. Gause GF. Experimental Analysis of Vito Volterra’s Mathematical Theory of the Struggle for Existence. Science. 1934;79(2036):16–7.

58. Schoener TW. Resource partitioning in ecological communities. Science. 1974;185(4145):27–39.

59. Mendes-Soares H, Krishnan V, Settles ML, Ravel J, Brown CJ, Forney LJ. Fine-scale analysis of 16S rRNA sequences reveals a high level of taxonomic diversity among vaginal Atopobium spp. Pathog Dis. 2015;73(4).

60. Yildirim S, Yeoman CJ, Janga SC, Thomas SM, Ho M, Leigh SR, et al. Primate vaginal microbiomes exhibit species specificity without universal Lactobacillus dominance. ISME J. 2014;8(12):2431–44.

61. Stumpf RM, Wilson BA, Rivera A, Yildirim S, Yeoman CJ, Polk JD, et al. The primate vaginal microbiome: comparative context and implications for human health and disease. Am J Phys Anthropol. 2013;152 Suppl 57:119–34.

62. Rivera AJ, Stumpf RM, Wilson B, Leigh S, Salyers AA. Baboon vaginal microbiota: an overlooked aspect of primate physiology. Am J Primatol. 2010;72(6):467–74.

63. Duar RM, Lin XB, Zheng J, Martino ME, Grenier T, Perez-Munoz ME, et al. Lifestyles in transition: evolution and natural history of the genus Lactobacillus. FEMS Microbiol Rev. 2017;41(Supp_1):S27–S48.

64. Mendes-Soares H, Suzuki H, Hickey RJ, Forney LJ. Comparative functional genomics of Lactobacillus spp. reveals possible mechanisms for specialization of vaginal lactobacilli to their environment. J Bacteriol. 2014;196(7):1458–70.

65. Lazaridis I. The evolutionary history of human populations in Europe. Curr Opin Genet Dev. 2018;53:21–21.

66. Serrano MG, Parikh HI, Brooks JP, Edwards DJ, Arodz TJ, Edupuganti L, et al. Racioethnic diversity in the dynamics of the vaginal microbiome during pregnancy. Nat Med. 2019;25(6):1001–11.

67. Greenbaum S, Greenbaum G, Moran-Gilad J, Weintraub AY. Ecological dynamics of the vaginal microbiome in relation to health and disease. Am J Obstet Gynecol. 2019;220(4):324–35.

